# Discovery of COVID-19 Inhibitors Targeting the SARS-CoV2 Nsp13 Helicase

**DOI:** 10.1101/2020.08.09.243246

**Authors:** Mark Andrew White, Wei Lin, Xiaodong Cheng

## Abstract

The raging COVID-19 pandemic caused by SARS-CoV2 has infected millions of people and killed several hundred thousand patients worldwide. Currently, there are no effective drugs or vaccines available for treating coronavirus infections. In this study, we have focused on the SARS-CoV2 helicase (Nsp13), which is critical for viral replication and the most conserved non-structural protein within the coronavirus family. Using homology modeling and molecular dynamics approaches, we generated structural models of the SARS-CoV2 helicase in its apo- and ATP/RNA-bound conformations. We performed virtual screening of ~970,000 chemical compounds against the ATP binding site to identify potential inhibitors. Herein, we report docking hits of approved human drugs targeting the ATP binding site. Importantly, two of our top drug hits have significant activity in inhibiting purified recombinant SARS-CoV-2 helicase, providing hope that these drugs can be potentially repurposed for the treatment of COVID-19.

## INTRODUCTION

A novel strain of severe acute respiratory syndrome coronavirus-2 (SARS-CoV2) is responsible for the current COVID-19 pandemic.^1,2^ Coronaviruses (CoVs) are enveloped 5’-capped, polyadenylated, single-stranded non-segmented, positive sense RNA viruses that cause various diseases in animals.^3^ In humans, manifestations of CoV infection range from asymptomatic, common cold, to lethal viral respiratory illness.^4^ There are no effective drugs or vaccines to treat or prevent CoV infection. Therefore, developing novel therapeutics for CoV represents an urgent medical need to combat the current COVID-19 devastation.

Upon infecting host cells, CoVs assembles a multi-subunit RNA-synthesis complex of viral non-structural proteins (Nsp) responsible for the replication and transcription of the viral genome.^4^ Among the 16 known CoV Nsp proteins, the Nsp13 helicase is a critical component for viral replication and shares the highest sequence conservation across the CoV family, highlighting their importance for viral viability. As such, this vital enzyme represents a promising target for anti-CoV drug development.^5–7^

To date, there is no atomic structure of SARS-CoV2 Nsp13 available, and none of the existing structural homologues (Table S1) published are suitable for molecular docking analyses. The two available apo-Nsp13 crystal structures are from the SARS-CoV (6JYT)^8^ and MERS-CoV (5WWP).^9^ Both 6JYT and 5WWP contain two identical chains in their crystal lattice: S1A and S1B, or M1A and M1B, respectively. The major difference between the two Nsp13 structures is associated with the 333-353 loop of the Rec1A^10^ domain that interacts with domain 1B, which is absent in M1A due to it being highly dynamic. The RMSD between M1B and S1A decreases from 1.57Å to 0.64Å when excluding this loop (Table S1). M1A and M1B have a larger difference in their Rec1A-Rec2A orientations than that among S1A, S1B and M1B. The CH and Stalk domains are similar (RMSDs<1 Å), while the orientations of the nucleotide binding domains (Rec1A and Rec2A) vary relative to them. The domain 1B among S1A, S1B and M1B are similar except for loops 202-208 which interacts with the Rec2A domain. The Rec2A domains are similar, except in the C-Terminus, and several flexible loops. The Rec2A domain seems to be intrinsically flexible with the crystallographic, intra-species, A/B domains having larger RMSDs than the inter-species Rec2A domains (Table S1). These apparent structural dynamics of Nsp13 structures highlight the value of having templates of the highly flexible helicase in multiple conformations, one of which may be a better target for high-affinity inhibitors. Therefore, we generated a series of SARS-CoV2 Nsp13 homology models in its apo- or substrate-bound states and performed *in-silico* docking, high-throughput virtual screening (HTvS), using all these models to search for potential SARS-CoV2 inhibitors.

## RESULTS AND DISCUSSION

### Homology model of SARS-CoV2 Nsp13 apo-structure based on SARS-CoV Nsp13 (6JYT)

The SARS-CoV2 Nsp13 helicase shares a 99.8% sequence identity to SARS-CoV (SARS) Nsp13 helicase with only one single residue difference (Figure 1a). The SARS Nsp13 helicase crystal structure was solved in its apo-state at a reported resolution of 2.8 Å.^8^ The crystallographic asymmetric unit contains two Nsp13 chains (S1A and S1B), offering a glimpse at the intrinsic flexibility of this helicase (Table S1). This crystal structure would be an ideal candidate for virtual screening, since as a homology model it differs from the SARS-CoV2 Nsp13 in only one amino acid, I570V, which is located away from the ATP and RNA binding sites. However, a close examination of the ATP binding site in this structure found several problems. First, the published model and electron density are of significantly lower quality than the MERS-CoV Nsp13 structure.^9^ The MERS structure was used to fill-in gaps in the SARS Nsp13 model, such as the dynamic 1B domain, which they were not able to model in their SAD-phased maps. In addition, the Walker-A loop,^10^ (residues G282-G287, K288, S289), which is responsible for binding the ATP substrate’s phosphates, has poor fit to the density and was built into the electron density with a highly improbable cis-peptide. Most significantly, the SARS-CoV2 Nsp13 structure does not have the usual sulfate ions bound even though there was clear difference density for the missing sulfate ions (Figure S1a). These errors in the published SARS Nsp13 structure made it unsuitable for use in molecular docking. Therefore, we decided to correct these errors and crystallographically re-refine the structure. On the other hand, the published apo crystal structure of MERS Nsp13, which retains 72% identity to the SARS-CoV2 Nsp13 helicase, has a better defined density around the Walker-A loop and sulfate ions (Figure S1b).^9^ Hence, we rebuilt the Walker-A loops in both chains guided by the MERS-Nsp13 structure, and added missing SO_4_ ions as supported by the difference electron density. In addition, we fixed other stereochemistry issues using the Crystallographic Object-Oriented Toolkit (COOT)^11^ and refined the structure in Phenix,^12^ following standard crystallographic procedures. This SARS Nsp13 template resulted in two models of apo-SARS-CoV2 Nsp13, S2A and S2B (Table S2).

**Figure 1.**
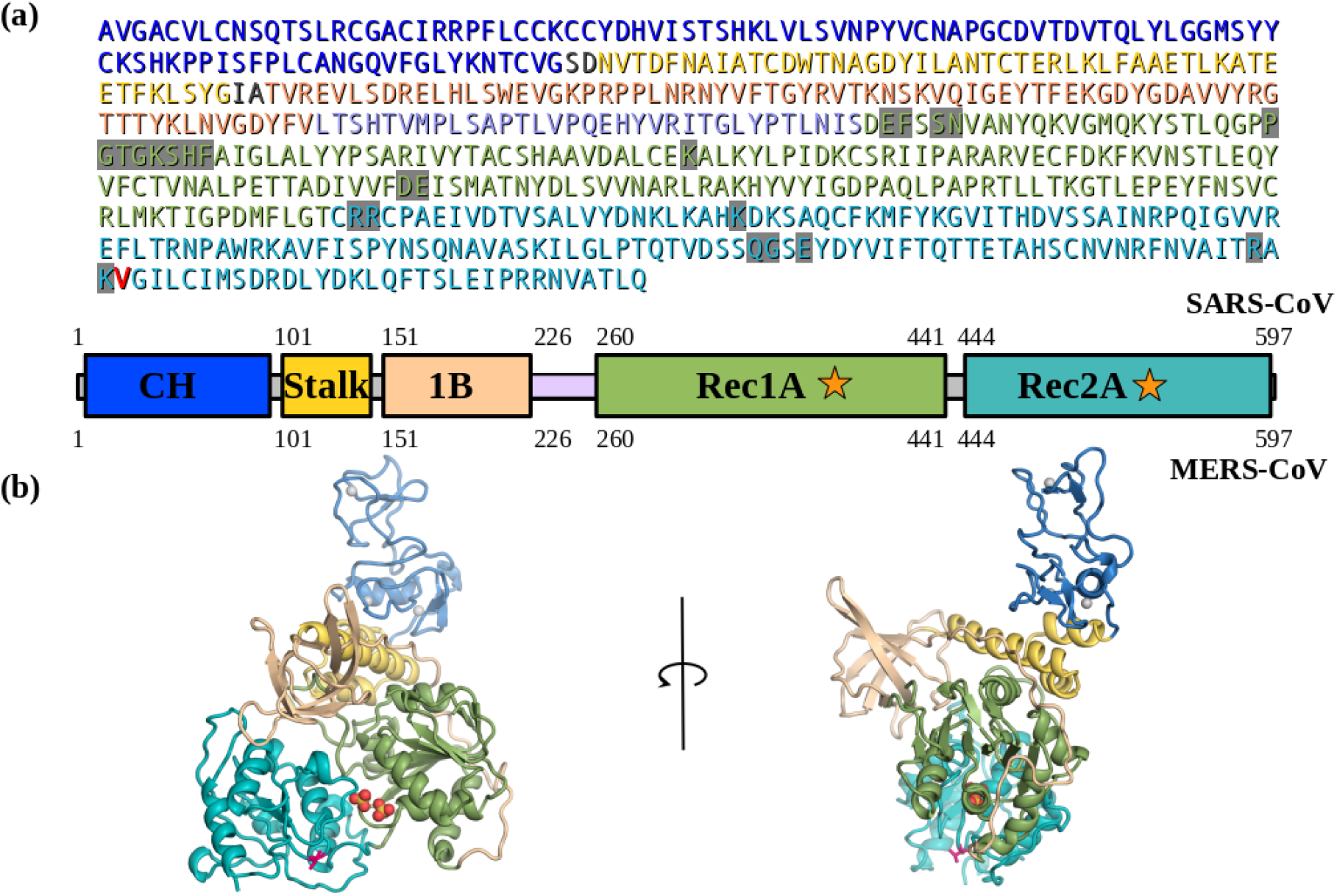
The SARS-CoV2 corona virus’s Nsp13 helicase structure. (a) Sequence and domain structure of SARS-CoV2. The ATP binding site residues are highlighted in grey. The single residue V570 that is different between SARS-CoV2 and SARS (I570) in the Rec2A domain, is coloured red. The domain structure and colouring is shown below the sequence. (b) The (apo) SARS-CoV2 Nsp13 structural model (S2A) based on the I570V mutation of SARS Nsp13 (6JYT), coloured-by-domain, the V570 is show as red sticks. The domain structure and colouring scheme are the same as shown above.

### The apo-SARS-CoV2 Nsp13 model based on MERS Nsp13 (5WWP)

The published MERS Nsp13 crystal structure,^9^ also consisting of two molecules (M1A and M1B) in the asymmetric unit (Table S1), was first refined using Phenix and COOT. We build the SARS-CoV2 homology models using this refined MERS crystal structure as a template, mutating the protein sequence. The resulting SARS-CoV2 models were then energy minimized by maintaining the crystallographic orientation for common atoms using Phenix, while stereochemical issues caused by mutation were corrected using COOT. The two homologous structural models based on either 6JYT (S2A and S2B) or 5WWP (M2A and M2B) are highly similar as expected with the Rec1A & Rec2A domains having a Cα-RMSD of only 1.4 Å (Figure S2, Table S2). The major changes being in the dynamic 1B domain, a slight rotation of the CH, Zn-binding, and the ATP-clamping Rec2A domains, plus a few flexible loops. The M2B model is the only complete apo-Nsp13 structure without gaps in the model.

### The SARS-CoV2 Nsp13:ATP:ssRNA complex model based on MERS Nsp13 and Upf1

The yeast Upf1 helicase complex^13^ with both ADP-AlF4 and ssRNA bound was previously identified as a structural homologue of apo MERS-CoV Nsp13 (5WWP) structure.^9^ While the yeast Upf1 helicase shares low sequence homology to CoV helicases (24% identity, 37% similarity), it has high structural homology to the SARS and MERS Nsp13 helicases, permitting the domains to be aligned by their secondary structure elements. Therefore, it represents a suitable template of the ATP and RNA-bound conformation of the CoV family helicases. We used the Upf1 complex structure to guide the modeling of SARS-CoV2 Nsp13/ATP/ssRNA complex structure (S2C) prior to energy minimization. In particular, modeling the domain motions upon complex formation (Figure 2, Table S2).

**Figure 2.**
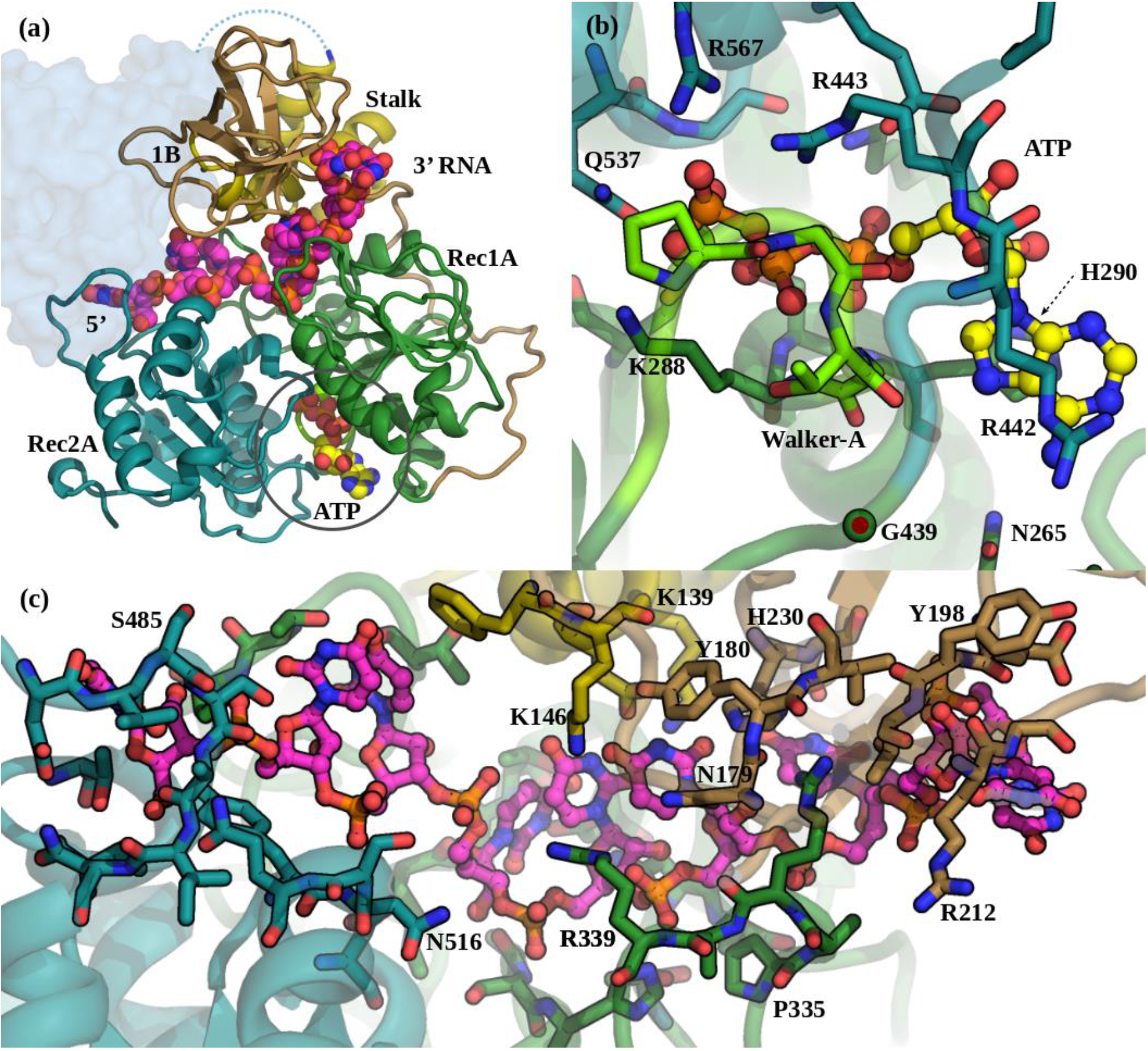
The SARS-CoV2 Nsp13:ATP:ssRNA complex model (S2C). (a) The Domain organization in the S2C complex. Colouring is as in Figure 1. The ATP carbons are coloured yellow and the ssRNA carbons are magenta, both shown as spheres. The missing CH, Zn-binding, domain is shown as a pale-blue blob. (b) The ATP binding pocket showing specific interactions with Nsp13. View is from below the Rec2A (cyan) towards the Rec1A (green) domain. The Walker-A loop is bright green. (c) The ssRNA binds between the Stalk, Rec1A, and Rec2A domains. View is from the Stalk (yellow) towards the Rec1A (green) - 1B (tan) domain interface.

Since the structural similarity of the CH Zn-binding domains of Upf1 and SARS-CoV2 is very low and the CH is not directly involved in substrate binding, we excluded it from the homology model of the Nsp13 complex. The alignment of the Nsp13 homology model, based on MERS Nsp13 chain B (M2B), to Upf1 started with the Rec1A domain, which includes the Walker-A loop. In this orientation the Stalk was also in good alignment. The two other domains, 1B and Rec2A both rotate into their substrate binding conformations, which moves them toward their substrates. The 1B domain has the largest motion to clamp against the 3’ end of the ssRNA in contact with the Stalk (Figure 2c, 5b). The Rec1A’s 333-350 loop, which was mostly disordered in the SARS crystal structures, shows motion upon ssRNA binding. The Rec1A Pro335-Arg337-Arg339 loop interacts with the RNA and forms a bridge with the Asn179-Tyr180 loop in domain 1B, which along with the Stalk domain’s terminal helix residues, Glu143 and Lys146, is also in contact with the RNA. The Rec1A domain forms the floor of the RNA-binding tunnel which has a mixture of back-bone amides, plus the Asn361, and His311 amines coordinating the backbone phosphates, as well as several carboxyl and hydroxyl groups hydrogen-bonding with the riboses and bases. The 478-490 loop of the Rec2A domain, moves out and away from the Stalk, increasing the space to accommodate the binding of the 5’ end of the ssRNA (Figure 2c).

For the ATP binding pocket, the Walker-A loop closes around the bound ATP, relative to the apo structures. The conventional A-loop adenine ring-distal tyrosine pi-stacking motif^14^ is replaced by a H290 in the helix immediately following the Walker-A, adjacent to the S289, which coordinates the Mg^2+^ ion (Figure 2b). The Rec2A domain rotates around the G439 link to Rec1A, bringing the Rec2A domain 5 Å closer to cover the ATP and also bringing Q537 into coordination with both the Mg^2+^ and the ATP-γP. The Rec2A’s arginine finger, R567, moves into the pocket to coordinate the ATP-γP, along with R443 which coordinates both the ATP-O3α and O3β (Figure 2b). At the exit of the ATP binding site, N265 and S264 are both poised to interact with the adenine ring’s NH2, while the adenine ring has hydrophobic stacking interactions with H290 and the alkane side-chain of R442. Together these features distinguish the ternary complex from the apo-Nsp13 models as a template for the active helicase.

### *In-silico* screening

HTvS was performed using the 5 homology models that we have generated. The ATP binding site for all four apo-models was identified by homology to the complex model (S2C). A suitable 30×26×26 Å^3^ box, centered on the ADP and encompassing the Rec1A Walker-A loop plus the opposing arginine finger in Rec2A,^10^ was defined and used for HTvS (Figure S4a). The Enamine Libraries (AC and PC) and ZINC-in-Trials Library totaling roughly 970,000 compounds were used for virtual screening of the ATP binding sites in the five models. For this study we focus on screening results based on the ZINC-in-Trials Library (9,270 compounds), which resulted in 369 drug hits approved for human use (Figure S3a). The detailed HTvS information for all 369 of these drugs from the ZINC Library is provided in the Excel spreadsheet (Table S3), part of the Supporting Information. Chemical affinity propagation clustering of the 369 HTvS selected drugs found 93 exemplar compounds highlighting the chemical diversity of the drugs selected by HTvS. These 369 drug hits can be filtered to a smaller number using a lower limit cut-off. For example a Total-Score cutoff of 80 will yield a list of 31 potential Nsp13 inhibitors for *in-vitro* or *in-vivo* screening assays (Figure S3a). The multiple targets produced groups of overlapping and non-overlapping hits (Figure S3b), which is to be expected since they include the ATP-bound and four apo-ATP sites, which were unrestrained by bound ligands.

### Top scoring apo-ATP site docking hits

The four models of the apo-Nsp13 helicase provided possible targets of the dynamic helicase. The top-scoring hits in these apo-Nsp13 ATP binding sites (Figure 3, Table S3) include: Cepharanthine^15^, Cefoperazone,^16^ Dihydroergotamine,^17^ Cefpiramide,^18^ Ergoloid (Dihydroergocristine (DHEC)),^19^ Ergotamine,^20^ Netupitant,^21^ Dpnh (NADH), Lifitegrast,^22^ Nilotinib,^23^ and Tubocurarin.^24^ The top-scoring hit Cepharanthine (CEP) is an anti-inflammatory drug used in Japan since the 1950s to treat a number of acute and chronic diseases, including the treatment of leukopenia and alopecia.

**Figure 3.**
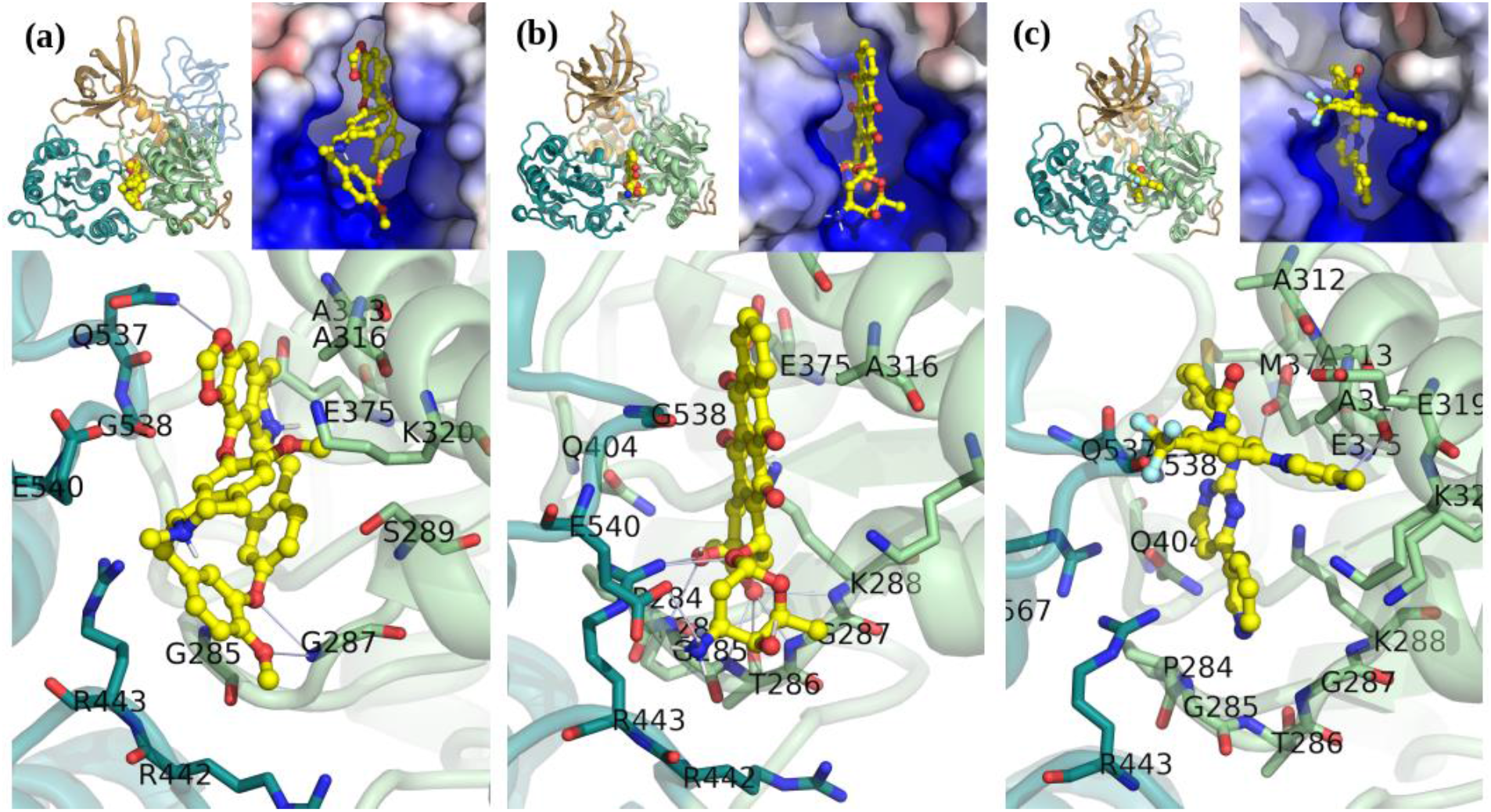
A Selection of apo-Nsp13 ATP binding site hits from virtual screening. (a) Cepharanthine shown fit in the (S2A) SARS Nsp13 structure’s ATP binding site, (b) Idarubicin in M2B, (c) Nilotinib in M2B. Inset: Left: Overview of the inhibitor bound to the Nsp13. Right: The electrostatic surface in the active site (colour gradient: blue-to-red +/-5 V).

### Top scoring complex-ATP site docking hits

In the complex the top scoring drugs were: Lumacaftor,^25^, Emend (Aprepitant),^26^ Nilotinib,^23^ Irinotecan,^27^ Enjuvia,^28^ Zelboraf,^29^ Cromolyn,^30^ Diosmin,^31^ Risperdal,^32^ and Differin (Adapalene).^33^ Surprisingly, although the ATP-binding pocket undergoes significant conformational changes upon ATP binding there is a significant overlap in hits even in this small group, with Lumacaftor, Nilotinib, Liftegrast, Idarubicin (Duanorubicin, Valrubicin), and Irinotecan scoring in the top-20 for both the apo- and ATP-bound models (Figure 4, Table S3).

**Figure 4.**
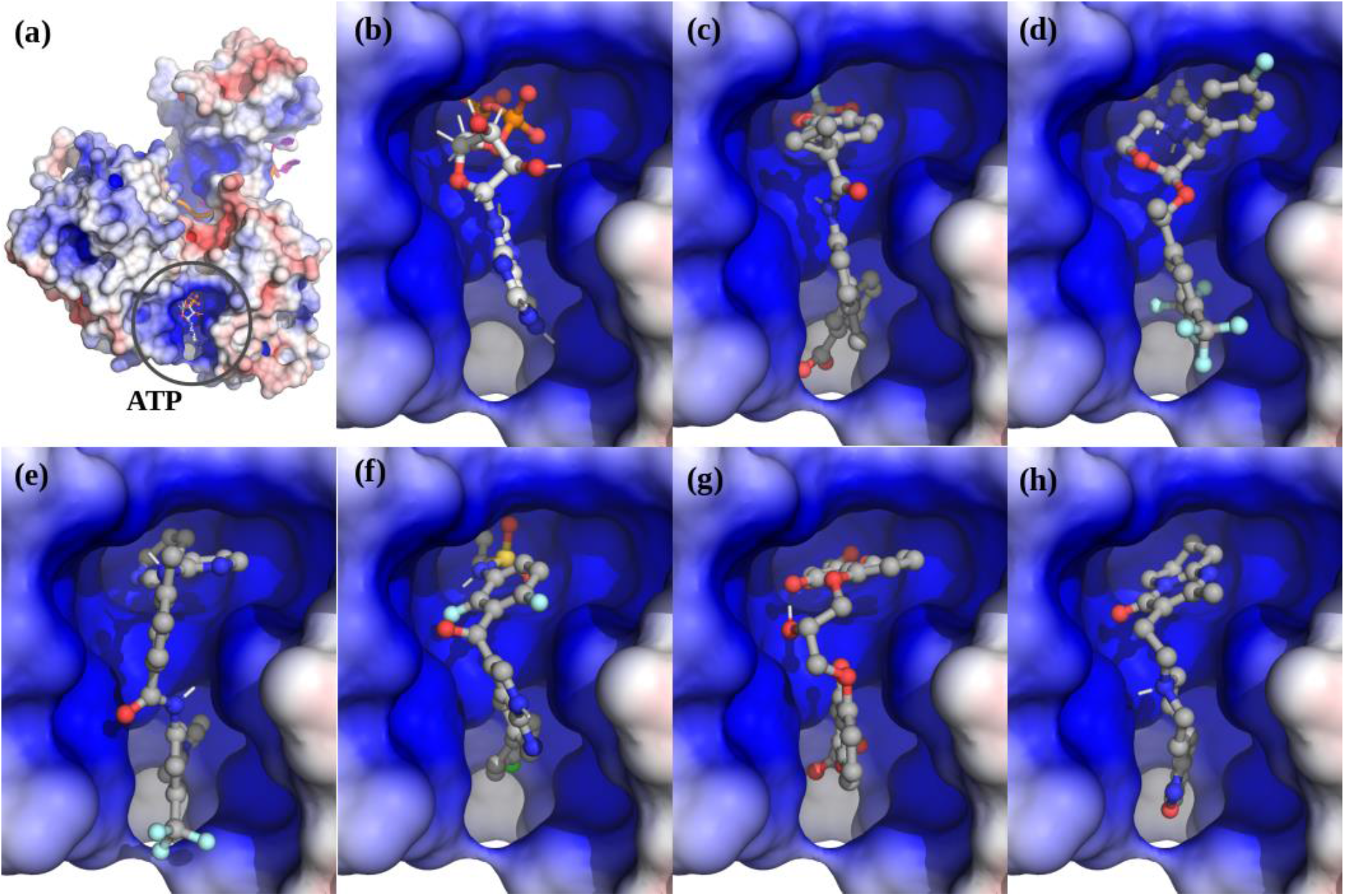
The top scoring hits for the Nsp13 complex ATP-site using virtual screening. (a) The Nsp13:ATP:RNA complex. (b) ATP. (c) Lumacaftor. (d) Emend (Aprepitant). Row-2: (e) Nilotinib. (f) Zelboraf. (g) Cromolyn. (h) Risperdal.

The two drugs Nilotinib^23^ and Lumacaftor^25^, were common top-scoring hits to the ATP site in both the apo- and ATP-bound models. Lumacaftor acts as a chaperone of protein folding and improves the processing of the most common cystic fibrosis transmembrane conductance regulator (CFTR) mutant, F508del, and its transport to the cell surface. Lumacaftor, in combination with Ivacaftor, is approved for the treatment of F508del CTTR.^34^ On the other hand, Nilotinib is a second generation Bcr-Abl tyrosine kinase inhibitor for the treatment of chronic myelogenous leukaemia (CML). Nilotinib was designed based upon the crystal structure of the Imatinib-Abl complex^35^ to fit into the ATP-binding site of the BCR-ABL protein with higher affinity.^36^ Nilotinib has also been identified as a potential inhibitor of the SARS-CoV-2 Nsp12-Nsp7-Nsp8 complex, which responsible for the RNA-dependent RNA polymerase activity, another key component of the multi-subunit RNA-synthesis complex.^37^

### Validation of HTvS hits

To validate our HTvS results, we ordered pure compound powder for several top hits and tested their ability to inhibit the ATPase activity of purified recombinant SARS-CoV-2 Nsp13 protein. Among the 10 drug candidates tested, two of our top HTvS hits Lumacaftor and Cepharanthine displayed activity in inhibiting Nsp13 ATPase activity with estimated IC50 values of 0.3 and 0.4 mM, respectively (Figure 5).

**Figure 5.**
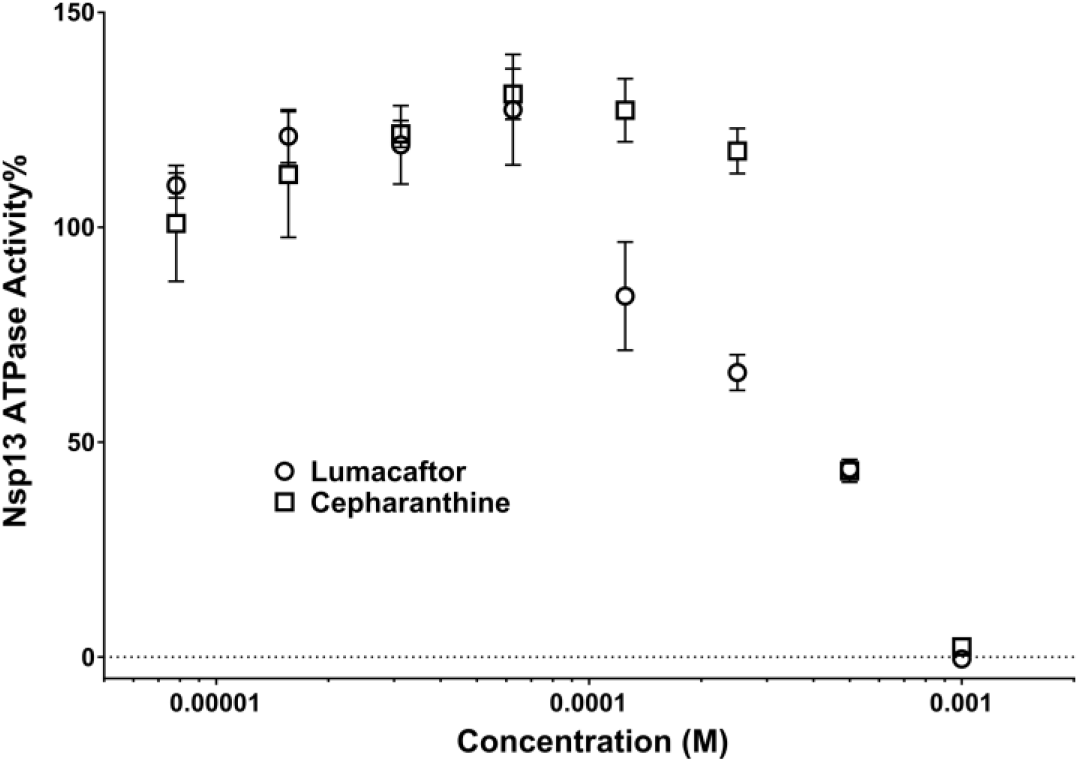
Inhibition of SARS-CoV-2 Nsp13 ATPase activity. Lumacaftor (o) and Cepharanthine (□). Data are presented as Mean ± SEM (N = 3).

Cepharanthine was the top scoring hit for our apo-ATP site screens while Lumacaftor was the top scoring hit for complex-ATP site screen (Figures 3 and 4, Table S4). Experimental confirmation of our top HTvS drug hits validates our structural models and HTvS approach targeting multiple conformational states. Cepharanthine is a strong candidate of potential anti-vials for the treatment of COVID-19.^38,39^ Cepharanthine was previously identified as an inhibitor of SARS.^40^ It has been found to effectively inhibit SARS-CoV2 in high throughput drug screening assay^41^ and in a separate siRNA assay,^42^ both using Vero cells. It has also been identified as a potential inhibitor of the SARS-CoV2 Nsp12-Nsp8-Nsp7 complex.^37^ Results from this study suggests that Cepharanthine may also synergistically target Nsp13 within the viral replication complex. On the other hand, Lumacaftor has not been directly implicated to play a role in inhibiting SARS-CoV2. Our study provides strong evidence to support the further testing of Lumacaftor in cell- and animal-based COVID-19 models.

## CONCLUSIONS

In conclusion, we have performed extensive integrated structural modeling to build atomic structural models of SARS-CoV2 Nsp13 in its apo- and substrate-bound conformations. Virtual molecular docking analyses targeting the ATP binding pocket using these structural models have led to the identification of potential inhibitor compounds, many of them approved human drugs. Of particular interest, two of our top HTvS hits show significant activity in inhibiting purified recombinant SARS-CoV-2 helicase, providing hope that these drugs can be potentially re-purposed for the treatment of COVID-19.

## EXPERIMENTAL METHODS

### Homology modeling of SARS-CoV2 Nsp13

Homology models of the SARS-CoV2 Nsp13 helicase were generated using the two available SARS^8^/MERS^9^ Nsp13 crystal structures. Issues we found with the highly similar SARS Nsp13 (6JYT) crystal structure prompted us to rebuild it in COOT^11^ and refine the new model in Phenix^12^ using standard crystallographic techniques (See Results, Figure S1). Mutation of the MERS (5WWP) crystal structure to the SARS-CoV2 sequence was performed in COOT. Energy minimization was performed in Phenix, followed by optimization of stereochemistry in COOT for several rounds.

### Molecular Dynamics

After rough domain alignment of the MERS-based apo SARS-CoV2 Nsp13 model to the Upf1 crystal structure, the domain linkers were remodeled in Coot with full stereochemical energy minimization, using the Upf1 electron density as a guide. Energy minimization of this SARS2-CoV-2:ATP:ssRNA complex model was performed using NAMD^43^, either through the VMD-NAMD^44^ interface or command line scripts. A 10ns MD run, with implicit water, was then performed, which permitted further motion of the domains. This model was then placed in an equilibrated TIP3 water box, with 0.15 mM NaCl, for further rounds of equilibration, annealing, and energy minimization, totaling 50 ns.

### HT Virtual Screening of the Models

Potential sites for inhibitor binding were identified by homology to the ATP or RNA binding sites in the structurally similar Upf1 helicase complex. We screened the identified substrate binding sites in each Nsp13 model using an implementation of AutoDock Vina^45^ on the Drug Discovery Portal at TACC.^46^ The ZINC^47,48^ drugs in Trials (9,270 compounds) Library, the Enamine-PC (84,359 compounds), and Enamine-AC (876,985 compounds) were used in high-throughput virtual screening of the five targets. Only the ZINC library “in-Trials” subset contains drugs currently approved for human use. Chemical clustering of hits was performed using the Affinity Propagation Clustering algorithm with Soergel (Tanimoto coefficient) distances on the ChemBio^49^ server (https://chembioserver.vi-seem.eu/).

### Expression and purification of recombinant SARS-CoV2 Nsp13 protein

The expression vector for SARS-CoV2 Nsp13 protein was constructed by site-directed mutagenesis (I570V) using the pET28a vector for SARS-CoV Nsp13 (kindly provided by Dr. Jian-dong Huang) as a tempelate.^50^ SARS-CoV-2 Nsp13 protein was expressed in BL21(DE3) cells and purified using a Ni-affinity column and a FPLC Superdex 200 Increase column as described previously.^8^

### Nsp13 ATPase activity assay

A modified ATPase assay was carried out by measuring phosphate release using a colorimetric method based on complexation with malachite green and molybdate (AM/MG reagent) using 96-well plates.^51,52^ Briefly, 20 μL reaction mixtures, containing 25 mM HEPES (pH 7.5), 50 mM NaCl, 5 mM MgCl_2_, 1 mM DTT, 0.25 mM ATP and 150 nM of Nsp13 in the presence or absence of various concentration of inhibitor, were incubated at 37 °C for 20 min. 80 μL of AM/AG dye solution was added into the reaction buffer and incubated at room temperature for 5 min. The production of phosphate was measured by monitoring the absorbance at 620 nm using a Molecular Devices FlexStation 3 Microplate Reader.

## AUTHOR INFORMATION

The authors declare no competing financial interests.

## ACKNOWLEDGMENT

We would like to thank Professor Jian-Dong Huang from The University of Hong Kong for providing the expression vector for SARS-CoV Nsp 13 helicase. This work is supported by grants from the National Institute of Health R35GM122536, R01AI111464, and in part, through a Sealy and Smith Foundation grant to the Sealy Center for Structural Biology and Molecular Biophysics. The authors acknowledge the Texas Advanced Computing Center (TACC) at The University of Texas at Austin for providing HPC resources that have contributed to the research results reported within this paper. (URL: http://www.tacc.utexas.edu). Special thanks to Drs. Stan Watowich (UTMB) and Joe Allen (TACC) for their help with the Drug Discovery Portal at the TACC.

