## Supplemental Tables 1-3 and Figures 1-5 for "Discovery of COVID-19 Inhibitors Targeting the SARS-CoV2 Nsp13 Helicase"

**Table S1. Comparison of Available Nsp13 Models/Templates.**

| | **SARS** | **MERS** | **Sc Upf1** |
|---|---|---|---|
| PDB id | 6JYT | 5WWP | 2XZL |
| **Resolution (Å)** | 2.8 | 3.0 | 2.4 |
| **Model(A) CC** | 0.85 | 0.89 | 0.90 |
| **RC outliers (%)** | 5.0 | 0.9 | 0.4 |
| **Rotomer Outliers (%)** | 0.8 | 4.7 | 1.4 |
| **Walker-A Site** | Empty? | (2) SO4 | ADP-$Al_4$ |
| **Cis-Peptides (%)[1]** | 0.80 | 0.48 | 0 |
| **Clash-score** | 29 | 15 | 8 |
| **#RSRZ>2 Percentile** | 1$^{st}$ | 82$^{nd}$ | 84$^{th}$ |
| **By-Chain Cα-RMSD** | **A / B** | **A / B** | **Sc Upf1** |
| **S1A (Å)** | 0 / 1.18 | 1.83 / 1.73 | 24 |
| **S1A[2] (Å)** | 0 / 1.28 | 1.77 / 1.70 | 21 |
| **S1A-CH (Å)** | 0 / 0.34 | 0.88 / 0.38 | 5 |
| **S1A-Stalk (Å)** | 0 / 0.19 | 0.38 / 0.38 | 5 |
| **S1A-1B (Å)** | 0 / 0.65 | na / | 21 |
| **S1A-Rec1A[3] (Å)** | 0 / 0.63 | 0.63 / 0.64 | 11 |
| **S1A-Rec2A (Å)** | 0 / 1.81 | 1.61 / 1.43 | 11 |
| **M1B (Å)** | 1.73 / 1.66 | 1.38 / 0 | 22 |
| **M1B[2] (Å)** | 1.70 / 1.55 | 0.98 / 0 | 18 |
| **M1B-CH (Å)** | 0.74 / 0.81 | 0.58 / 0 | 8 |
| **M1B-Stalk (Å)** | 0.38 / 0.39 | 0.50 / 0 | 10 |
| **M1B-1B (Å)** | 1.24 / 1.43 | na / 0 | 8 |
| **M1B-Rec1A[3] (Å)** | 0.64 / 0.77 | 0.43 / 0 | 14 |
| **M1B-Rec2A (Å)** | 1.43 / 1.39 | 1.65 / 0 | 4 |

[1] Number of non-proline Cis-peptides. [2] Excluding CH Zn-binding domain. [3] Excluding the 333-353 loop which interacts with domain 1B.

**Table S2. The Cα-RMSD (Å) values for the four apo-SARS-CoV2 Nsp13 models.**

| Nom\Template | SARS_2A | SARS_2B | MERS_2A | MERS_2B |
|---|---|---|---|---|
| **S2A (Å)** | 0 | 0.74 | 1.88 | 1.67 |
| **S2B (Å)** | 0.74 | 0 | 1.90 | 1.71 |
| **M2A (Å)** | 1.88 | 1.90 | 0 | 1.61 |
| **M2B (Å)** | 1.67 | 1.71 | 1.61 | 0 |

The SARS_CoV Nsp13 (6JYT) chains A and B templates were used to create models S2A and S2B respectively. The MERS_CoV Nsp13 (5WWP) chains A and B templates were used to create models M2A and M2B respectively.

**Table S3. The Top-Twenty Drug Hits for each SARS-CoV2 Nsp13 model.***

| NN | (apo) | DRUG_Name | (S2A) | DRUG_Name | (S2B) | DRUG_Name | (M2A) | DRUG_Name | (M2B) | DRUG_Name | (ATP) | DRUG_Name |
|---|---|---|---|---|---|---|---|---|---|---|---|---|
| 1 | **-10.3** | **Cepharanthine** | **-10.3** | **Cepharanthine** | **-10.1** | **Dihydroergotamine** | **-10.2** | **Cefoperazone** | -9.2 | Ledipasvir | **-11.1** | **Lumacaftor** |
| 2 | **-10.2** | **Cefoperazone** | **-10** | **Ergoloid** | **-10** | **Ergotamine** | **-10** | **Cefpiramide** | -9.1 | Idarubicin | **-10.6** | **Emend** |
| 3 | **-10.1** | **Dihydroergotamine** | **-10** | **Ergotamine** | **-10** | **Netupitant** | -9.9 | Dpnh | -9 | Daunorubicin | **-10.5** | **Nilotinib** |
| 4 | **-10** | **Cefpiramide** | -9.9 | Dihydroergotamine | -9.9 | Ergoloid | -9.9 | Lifitegrast | -9 | Dpnh | **-10.4** | **Irinotecan** |
| 5 | **-10** | **Ergoloid** | -9.7 | Irinotecan | -9.9 | Nilotinib | -9.8 | Raltegravir | *-8.9* | *Carindacillin* | **-10.3** | **Enjuvia** |
| 6 | **-10** | **Ergotamine** | -9.7 | Nilotinib | -9.9 | Tubocurarin | -9.7 | Daunorubicin | *-8.8* | *Cefonicid* | **-10.1** | **Zelboraf** |
| 7 | **-10** | **Netupitant** | -9.7 | Vumon | -9.8 | Idarubicin | -9.6 | Avodart | *-8.8* | *Cefpiramide* | **-10** | **Cromolyn** |
| 8 | -9.9 | Dpnh | -9.6 | Conivaptan | -9.8 | Oxytetracycline | -9.6 | Idarubicin | *-8.8* | *Diosmin* | **-10** | **Diosmin** |
| 9 | -9.9 | Lifitegrast | -9.6 | Fosaprepitant | -9.7 | Dihydroergotoxine | -9.6 | Nilotinib | *-8.8* | *Rutin* | -9.9 | Differin |
| 10 | -9.9 | Nilotinib | -9.4 | Biosone | -9.7 | Dpnh | -9.5 | Carindacillin | *-8.7* | *Azelnidipine* | -9.9 | Risperdal |
| 11 | -9.9 | Tubocurarin | -9.4 | Ceftobiprole | -9.7 | Metocurine | -9.5 | Valrubicin | *-8.7* | *Carindacillin* | -9.9 | Rutin |
| 12 | -9.8 | Idarubicin | -9.4 | Venetoclax | -9.7 | Otc | -9.4 | Delamanid | *-8.7* | *Demeclocycline* | -9.9 | Valrubicin |
| 13 | -9.8 | Oxytetracycline | -9.3 | Ponatinib | -9.7 | Venetoclax | -9.4 | Ellence | *-8.7* | *Lumacaftor* | -9.8 | Pemetrexed |
| 14 | -9.8 | Raltegravir | -9.2 | Avodart | -9.6 | Daunorubicin | -9.4 | Exjade | *-8.7* | *Meclocycline* | -9.7 | Dihydroergotamine |
| 15 | -9.7 | Daunorubicin | -9.2 | Biosone | -9.6 | Doxycycline | -9.4 | Fosaprepitant | *-8.6* | *Cefoperazone* | -9.7 | Ergotamine |
| 16 | -9.7 | Dihydroergotoxine | -9.2 | Bromocriptine | -9.6 | Imatinib | -9.3 | Carindacillin | *-8.6* | *Doxycycline* | -9.7 | Idarubicin |
| 17 | -9.7 | Irinotecan | -9.2 | Cialis | -9.6 | Lumacaftor | -9.3 | Ceftobiprole | *-8.6* | *Methacycline* | -9.7 | Lifitegrast |
| 18 | -9.7 | Metocurine | -9.2 | Rutin | -9.6 | Meclocycline | -9.3 | Dolutegravir | *-8.6* | *Raltegravir* | -9.6 | Amaryl |
| 19 | -9.7 | Otc | -9.2 | Saquinavir | -9.6 | Methacycline | -9.3 | Doxorubicin | *-8.6* | *Sultamicillin* | -9.6 | Avodart |
| 20 | -9.7 | Venetoclax | -9.2 | Tasosartan | -9.5 | Biosone | -9.3 | Erismodegib | *-8.6* | *Tasosartan* | -9.6 | Ellence |

*The (apo) column is the highest scoring hit for any of the apo-ATP sites. The SARS_CoV Nsp13 (6JYT) chains A and B templates were used to create models S2A and S2B respectively. The MERS_CoV Nsp13 (5WWP) chains A and B templates were used to create models M2A and M2B respectively. The apo-Nsp13 ATP sites were selected as targets. The Upf1 guided Nsp13:ATP:ssRNA (S2C) complex had both ATP and ssRNA sites targeted. Dpnh: Dihydronicotinamide Adenine Dinucleotide Disodium Salt Hydrate (NADH). Cardinacillin is in the database as different stereoisomers (+/-). Otc: an isomer of oxytetracycline. The Vina docking scores are given in the column preceding the drug name. The column header specifies the target.

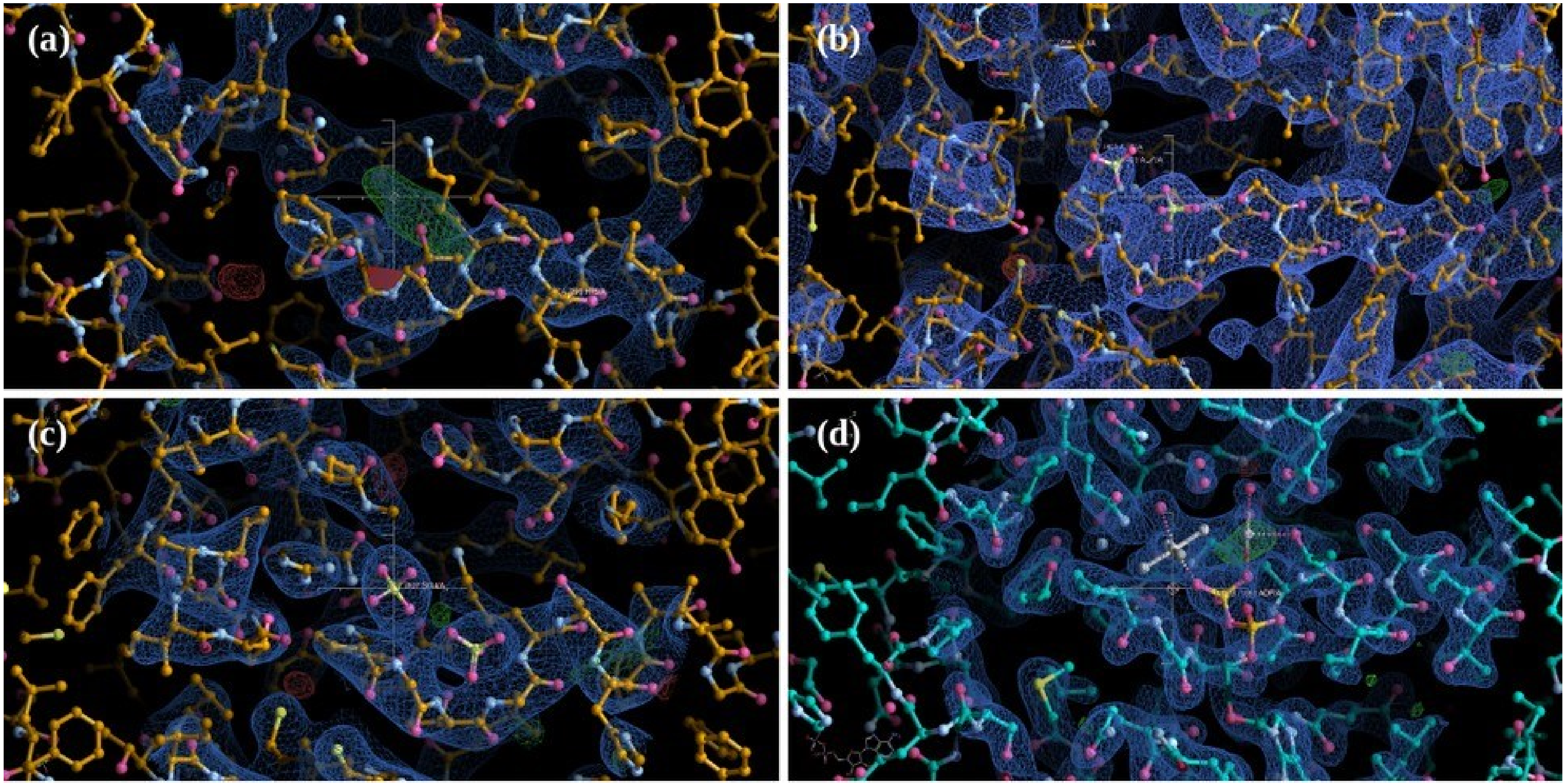


**Figure S1. The electron density of the Walker-A site in each structure.** (a) The SARS Nsp13 (6JYT) structure before and (b) after rebuilding the NTP site. (c) The MERS Nsp13 (5WWP) NTP binding site with two SO4 ions. (d) The yeast Upf1 (2XZL) ADP-AlF4 bound NTP site used as a template for the Nsp13 complex model.

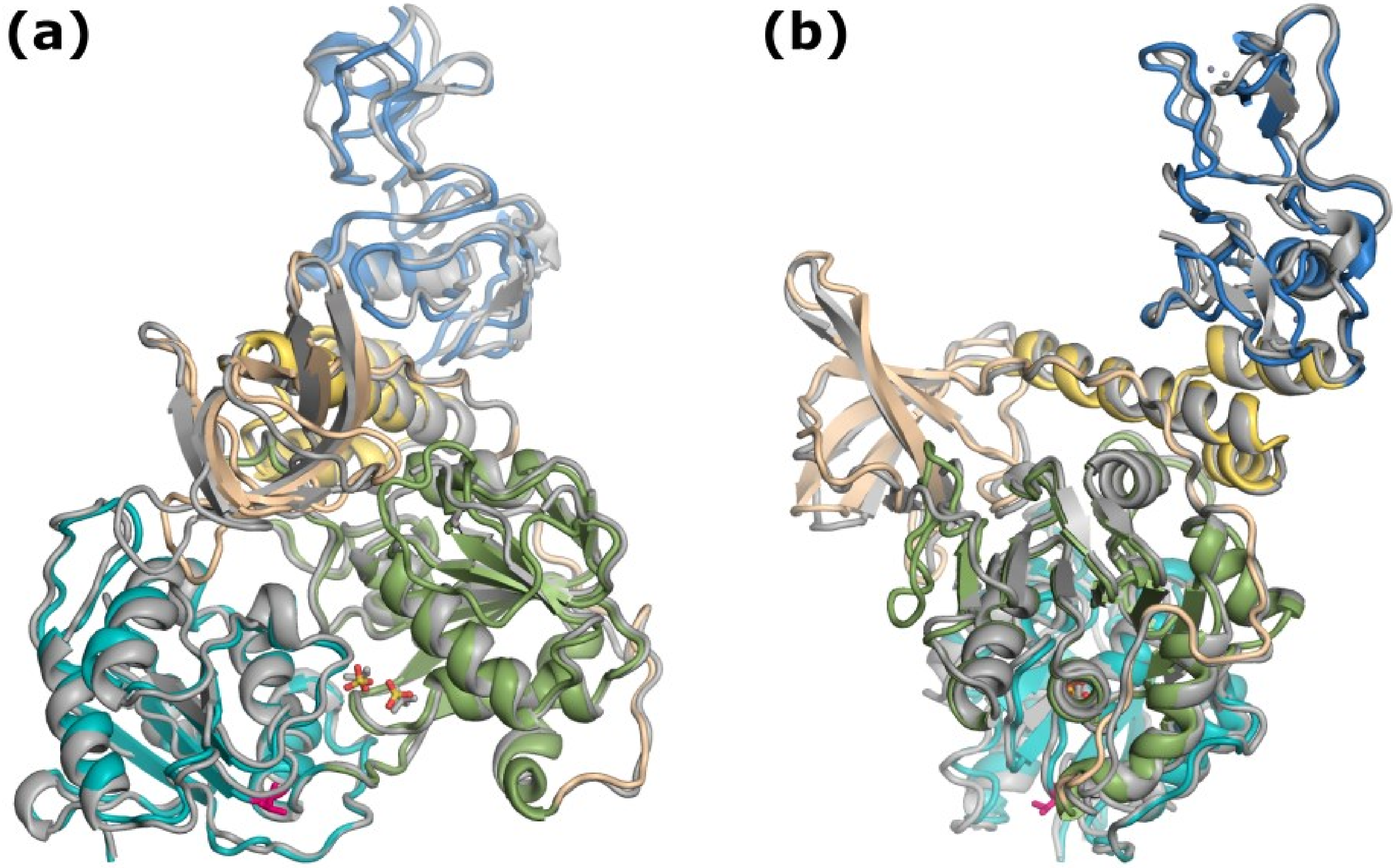


**S2. Comparison of the apo-Nsp13 Models.** Superposition of the (M2B) apo-SARS-CoV2 Nsp13 homology model based on MERS Nsp13 helicase (5WWP), coloured-by-domain as in Figure 1, and the (S2A) apo-SARS-CoV2 Nsp13 structural model (grey) based on the SARS Nsp13 (6JYT) with the I570V mutation, show as red sticks. The two views (a and b) are rotated by 90° about the vertical. The Walker-A site is occupied by two $SO_4$ ions, lower center, in the apo crystal structures. See Figure S5 for superposition of other apo-Nsp13 models.

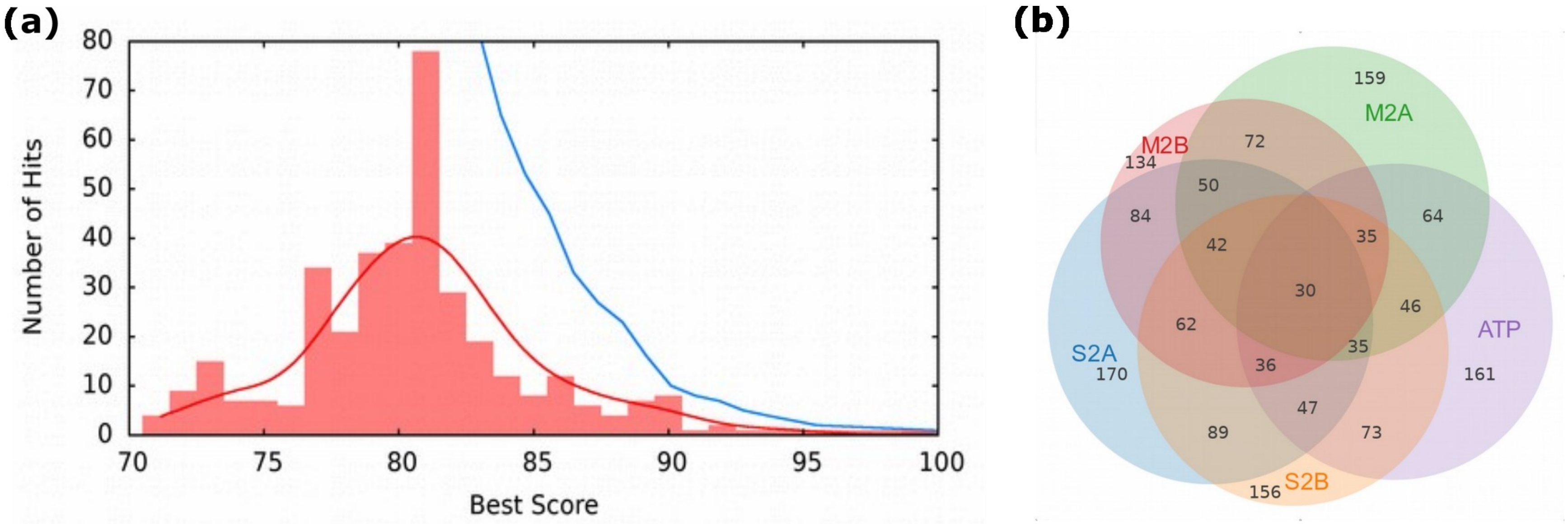


**Figure S3. Selected top hits.** (a) Histogram of the 369 drugs discovered using *in-silico* screening of the SARS-CoV2 Nsp13 helicase models (Table S4). The best score (x-axis) is the best normalized Vina docking score for any model, higher is better. The red line is the k-density smoothed distribution. The cut-off filtered results (blue line) is the cumulative number of drugs as the cut-off is lowered. (b) The Venn diagram of the ATP binding site hits for the five ATP templates. Each intersection has the hit count number.

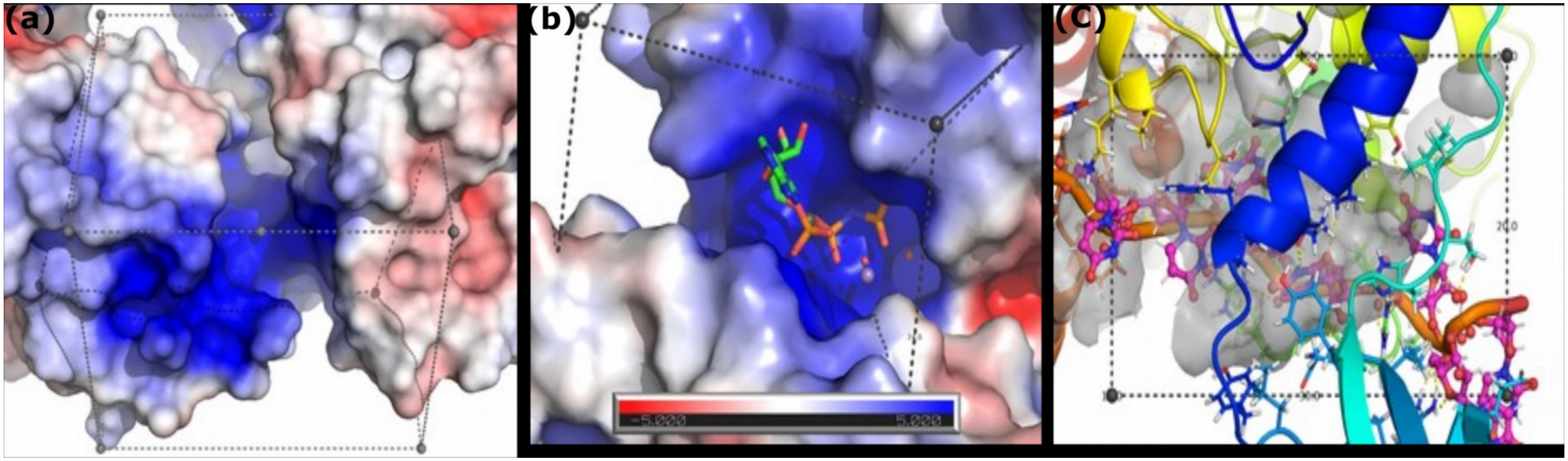


**Figure S4. SARS-CoV2 models' ATP-binding Site for HTvS.** Views of the (a) SARS-CoV (S2A), (b) MERS (M2B), and (c) Upf1 (S2C-ATP) based models of the SARS-CoV2 NTP binding site. ATP is modeled into a,b based on the Upf1 structure.

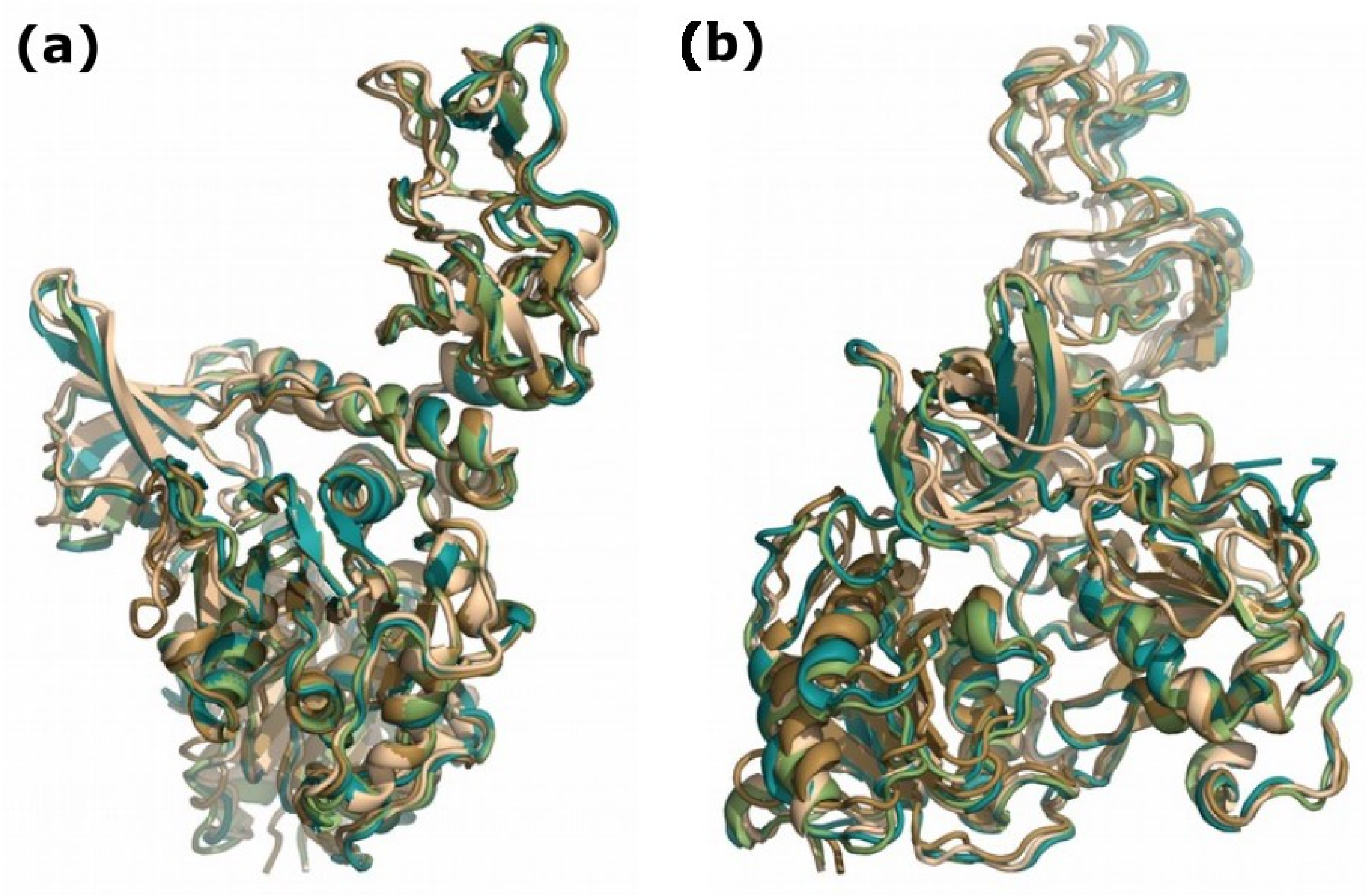


**Figure S5. Superposition of the four (apo) SARS-CoV2 Nsp13 homology models**: S2A (cyan), S2B (green), M2A (brown), M2B (tan). Inter-chain Cα-RMSDs are 1.6 Å for MERS, and 0.74 Å for SARS. The inter-species model Cα-RMSDs range from 1.7 to 1.9 Å (Table S-2). The two views (a and b) are rotated by 90° about the vertical.
